# Aphid transmission of a Potexvirus, *Foxtail mosaic virus*, in the presence of the Potyvirus helper component proteinase

**DOI:** 10.1101/2021.09.05.459040

**Authors:** Jun Jiang, Eric Yu, Clare L. Casteel

**Author notes:** Author for correspondence: *Clare L. Casteel*,.

## Abstract

To establish successful infections, plant viruses compete with the host plants for limited resources and thus alter the physiological state of the plants. After successful infection, insect vectors are required for the transmission of some plant viruses to the next host plant. One of the largest groups of plant viruses, the *potyvirus*, can be transmitted by aphids. During transmission, the potyvirus protein helper component proteinase (HC-Pro) binds to the yet-to-be-defined aphid receptor on the stylet, as well as to the virus particles through the Asp-Ala-Gly (DAG) motif of the viral coat protein. Previously it was determined that a naturally occurring DAG motif in the non-aphid transmissible *potexvirus*, *Potato aucuba mosaic potexvirus* (PAMV), is functional when the HC-Pro is provided through co-infection with a potyvirus. Further, the DAG motif of PAMV can be successfully transferred to another non-aphid transmissible potexvirus, *Potato virus X* (PVX), to convey aphid transmission capabilities. We expand on this previous work by demonstrating, the DAG motif from two different potyviruses, *Sugarcane mosaic virus* and *Turnip mosaic virus*, as well as the DAG motif from the previous potexvirus PAMV, can be added to another non-aphid transmissible *potexvirus*, *Foxtail mosaic virus (FoMV)*, to make it aphid transmissible. Transmission efficiency varied from less than 10% to over 80% depending on the DAG motif and host plant used in transmission, suggesting not all DAG motifs are equal for engineering aphid transmission. The underlying mechanisms mediating this variation still need to be explored.

## 1. Introduction

Plant viruses must take over the metabolism of the host to obtain resources and establish a successful viral infection. Interference with cellular homeostasis causes a variety of pathogenic effects, including stunting, discoloration, and malformation (Gergerich and Dolja, 2006). Many plant viruses move among plant hosts and plant populations through insect vectors. The rapid reproduction of some vectors and their ability to travel over long distance have led to the fast spread of plant viruses in ecosystems and significant amounts of economic loss. Fortunately, the transmission of plant viruses by insect vectors requires specificity, with a particular plant virus transmitted by a particular insect species in general (Casteel and Falk, 2016; Whitfield et al., 2015).

Depending on the virus transmission mode, insect vector-mediated plant virus transmission can be classified as persistent, semi-persistent, or non-persistent transmission. For persistent transmission, the plant viruses enter the insect body and may or may not replicate within the insect. For example, the *Rice dwarf virus* (RDV) enters the alimentary canal, further reaches the midgut of its leafhopper vector, and can multiply in the insect host (Chen et al., 2011b). On the contrary, the *Barley yellow dwarf virus* (BYDV) circulates in the aphid but does not replicate in this insect vector (Li et al., 2001). In these cases, the insect vectors are viruliferous for their entire life cycle. Other viruses, such as the *Lettuce infectious yellows virus* (LIYV), can be transmitted by their insect vectors in a non-circulative, semi-persistent manner, where the virions are retained for hours to days in the insect anterior foregut, or cibarium, and are lost after the insect molts (Chen et al., 2011a; Tian et al., 1999). In contrast, some plant viruses can be transmitted by their insect vectors in a non-persistent manner where they are only retained for minutes to hours in the insect stylet and are lost after the insect molts. For example, *Cauliflower mosaic virus* (CaMV), *Cucumber mosaic virus* (CMV), and *Potato virus Y* (PVY) are transmitted in a non-persistent manner by aphids, the primary insect vector for this transmission mode (Nanayakkara et al., 2012; Severin H, 1948; Swenson and Marsh, 1967).

The underlying molecular mechanism of non-persistent plant virus transmission by aphids has been studied extensively from the perspective of viral components through the purification of virus particles, the generation of mutated viruses, and the purification of transmission-related viral proteins. For example, the viral coat protein (CP) is the major determinant of aphid-mediated CMV transmission, especially the negatively charged loop structure on the surface of the virus particle which is critical for aphid transmissibility (Liu et al., 2002). The transmission of CaMV requires multiple viral proteins, of which the viral protein P2 binds to the aphid stylet through its N-terminus, and the C-terminus of P2 interacts with the virion-associated viral protein P3, for aphid-mediated transmission (Blanc et al., 2014; Hoh et al., 2010; Plisson et al., 2005). The transmission of potyviruses, including *Turnip mosaic virus* (TuMV), *Sugarcane mosaic virus* (SCMV), and PVY, is mediated by the Asp-Ala-Gly (DAG) motif (or the related variants, such as DTG, DAE, and DAA motif) of the CP and the Lys-Ile-Thr-Cys (KITC) and Pro-Thr-Lys (PTK) motif of the viral helper component proteinase (HC-Pro) (Blanc et al., 1998; Gadhave et al., 2020; Peng et al., 1998). In this case (and the previous with CaMV), a bridge model has been proposed: the viral protein HC-Pro simultaneously binds to the aphid stylet and the virus particle for the transmission of the virus. Many other viral proteins are dispensable for vector-mediated virus transmission, but they can facilitate virus transmission by modulating plant physiology and insect vector behavior and biology. For instance, the potyvirus NIa-Pro can disrupt the ethylene response of the host plant, thus attracting the insect vectors to the virus-infected plant (Bak et al., 2019; Casteel et al., 2015). The 2b protein of CMV also can alter the emission of volatile compounds, as well as enhance the production of reactive oxygen species, to promote the virus transmission by the aphid vectors (Guo et al., 2019; Tungadi et al., 2017). However, the aphid receptors involved in the virus transmission are still largely unknown, thus far only a few potential candidates have been discovered (Deshoux et al., 2020; Yang et al., 2008).

Potexviruses are not aphid transmissible, however, the CP from *Potato aucuba mosaic potexvirus* (PAMV) contains a DAG motif. While PAMV cannot be transmitted by aphids in a single infection (only PAMV), it can be transmitted by aphids when the potyviral HC-Pro is provided in mixed infections with a potyvirus (PVY) (Baulcombe et al., 1993). It was further demonstrated that aphid transmissibility can be successfully transferred to another potexvirus, *Potato virus x* (PVX), with the addition of PAMV’S DAG motif to the N-terminus of PVX’s CP (Baulcombe et al., 1993). To expand on this work, we evaluated if the DAG motifs from different potyviruses can be added to another potexvirus, *Foxtail mosaic virus* (FoMV), to convey aphid transmissibility. We compared the transmission efficiency of FoMV with different DAG motif additions to its CP, including the addition of 15 or 35 amino acid residues from the CP of two potyviruses (SCMV or TuMV), or the addition of 20 or 40 amino acid residues of PAMV’s CP. The addition of the 35 amino acid residues from the N-terminus of SCMV’s CP resulted in the highest aphid transmission efficiency (up to 75%) to maize plants (*Zea mays*), while transmission success was less than 20% for all other recombinant viruses to maize. Overall, aphid transmission was more successful to the model plant *Nicotiana benthamiana* compared to maize, where transmission efficiency varied from 30% to over 80%, depending on the N-terminus addition. These results demonstrate not all DAG motifs are equal for engineering aphid transmissibility in potexviruses.

## 2. Materials and methods

### 2.1. Potexvirus and potyvirus sequence comparison

The CP amino acid sequence of potexvirus, *Cymbidium mosaic potexvirus* (CymMV, accession number AAL12628), *Narcissus mosaic virus* (NMV, accession number NP_040782), *Pepino mosaic virus* (PepMV, accession number AIL23124), *Bamboo mosaic virus* (BaMV, accession number AOS51152), PVX (accession number AAA47181), PAMV (accession number AXL97636), FoMV (accession number AWT40560), and of the potyvirus, PVY (accession number ADH52720), *Soybean mosaic virus* (SMV, accession number QBB78854), *Tobacco etch virus* (TEV, accession number ABH10566), *Plum pox virus* (PPV, accession number ABU97781), *Pepper mottle virus* (PepMoV, accession number BAB91329), *Bean yellow mosaic virus* (BYMV, accession number ABM69144), *Potato virus A* (PVA, accession number NP_734368), TuMV (accession number ANW35618) and SCMV (accession number P32652), were aligned with ClustalW2 online server (https://www.ebi.ac.uk/Tools/msa/clustalo/).

### 2.2. Molecular cloning

The FoMV infectious clone pCambia1380/FoMV was a generous gift from Professor Steven A. Whitham (Iowa State University, USA), and the TuMV infectious clone pCambia0380/TuMV:6K2-GFP (Cotton et al., 2009) was kindly provided by Professor Jean-François Laliberté (INRS-Institut Armand-Frappier, Canada). To make the mutants pCambia1380/FoMV^V8A-T9G^, pCambia1380/FoMV^T12G^, pCambia1380/FoMV^Y14A-K15G^, pCambia1380/FoMV^V8A-T9G-T12G^, and pCambia1380/FoMV^A6V-V8A-T9G^, three-step cloning processes were performed. First, the HSSB fragment, of which the *<u>S</u>ph*I-*<u>S</u>ac*II fragment of wild type (WT) FoMV that contains the N-terminal part of the FoMV CP coding sequence, with the addition of *<u>H</u>indIII* restriction site at the 5’ end and *<u>B</u>am*HI restriction site at the 3’ end, was cloned into the smaller vector pBluescript SK(+), and the resulting clone was pSK-HSSB. Site-directed mutagenesis was then performed, and the corresponding mutations were then introduced into pSK-HSSB using a complementary primer set. The *<u>S</u>ph*I-*<u>S</u>ac*II fragment of pSK-HSSB with the desired mutations was then subcloned into pCambia1380/FoMV.

To construct the FoMV mutants (pCambia1380/FoMV^SCMV N15^, pCambia1380/FoMV^TuMV N15^, pCambia1380/FoMV^PAMV N20^, pCambia1380/FoMV^SCMV N35^, pCambia1380/FoMV^TuMV N35^ and pCambia1380/FoMV^PAMV N40^), three-fragment Gibson assembly reactions were performed. In general, fragment 1 was PCR amplified with Primer F1 and R1, using the plasmid pCambia1380/FoMV as the template. This fragment was approximate 200 bp and located right upstream of the FoMV CP coding sequences. The coding sequences of SCMV N15, TuMV N15, PAMV N20, SCMV N35, TuMV N35, and PAMV N40 were incorporated into the primer R1. Fragment 2 was about 700 bp and was PCR amplified with primer F2 and R2 using the plasmid pCambia1380/FoMV as the template. This fragment contained mostly the FoMV CP coding sequences. Fragment 3 was the vector backbone of pCambia1380/FoMV that was digested with *Bsu36I* and *XbaI*. The primer sequences are listed in Supplemental Table 1. All constructs were confirmed by DNA sequencing.

### 2.3. Virus infection

All FoMV derived constructs were transformed into *Agrobacterium tumefaciens* (strain GV3101) and the positive transformants were selected on LB Kanamycin-Rifampicin agar plates. The positive transformants were cultured overnight, centrifuged, and suspended in a 10 mM MgCl_2_ and 150 μM acetosyringone solution. For viral RNA accumulation assays, the OD_600_ of *A. tumefaciens* suspensions was adjusted to 0.03; And to purify the virus particles for transmission electron microscopy (TEM) observations, the OD_600_ of *A. tumefaciens* suspensions was adjusted to 0.2. To co-infect *N. benthamiana* plants with FoMV recombinant virus and TuMV/6K2:GFP, the OD_600_ was 0.2 and 0.05 for the FoMV recombinant virus and TuMV/6K2:GFP *A. tumefaciens* suspensions, respectively. Agroinfiltration was done with four-week-old *N. benthamiana*, and the agro suspensions were infiltrated into the underside of the leaves with a needleless syringe. To test the infectivity of the recombinant viruses in maize plant, the inoculum was prepared by grinding the FoMV mutant infected *N. benthamiana* leaf tissues in 50 mM potassium phosphate buffer, pH 7.0 (1 g tissue to 4 ml buffer), and rubbed onto 1-week-old maize plants with carborundum. All plants were grown at 24 °C, 16-h-light/ 8-h-dark photoperiod.

### 2.4. Virus purification and preparation for TEM observation

Systemic infected *N. benthamiana* leaf tissues were collected 2 weeks after agroinfiltration. Leaf tissues were then homogenized with the same amount (1 g tissue to 1 ml buffer) of buffer containing 0.1 M Tris-Citric acid (pH 8.0), 0.2%β-mercaptoethanol, and 0.01 M sodium thioglycolate. Triton X-100 was added gradually to a final concentration of 1% in a period of 15 min with constant stirring at 4 °C. Chloroform was added to a final concentration of 25%, and constant stirring at 4 °C for 30 min. The mixture was then centrifuged at 12,000x g for 15 min to separate the water phase and organic phase. PEG6000 was then added to the water phase, to a final concentration of 5% and constant stirring at 4 °C overnight. The mixture was then centrifuged at 10,000x g for 15 min, and the resulting pellet was resuspended with 300 ul of the extraction buffer described above. To observe the virions under TEM, 5 ul of purified virions was loaded onto the pretreated grid. The solution was then absorbed with a filter paper after 5 sec. The same step was repeated 5 times but with 5 ul 2% phosphotungstic acid solution. The grid was then air-dried for 2 min and was ready for TEM observation. TEM observation was done with a JEOL 2100F transmission electron microscope at 200 kV accelerating voltage.

### 2.5. Aphid transmission

All aphid colonies and plants were maintained in the growth chamber (24 °C, 16-h-light/ 8-h-dark photoperiod). The green peach aphid (*Myzus persicae*) colony was reared on *N. tabacum*. The aphid adults were collected and then starved for 4 h. The aphids were then allowed to acquire the virus from the infected source tissues for 5 min and then were transferred to healthy *N. benthamiana* plants (2-week-old) or sweet corn (*Zea mays* cv. Golden Bantam, 1-week-old). Pesticide was sprayed to kill the aphids 24 h later. Two to three weeks after aphid transmission, the newly emerged leaves were collected for RT-PCR.

### 2.6. RT-PCR and RT-qPCR Analysis

The agroinfiltrated leaf tissues were collected for RT-qPCR analysis, while either the systemic infected or newly emerged leaf tissues were collected for RT-PCR analysis. The total RNAs were isolated from the leaf tissues using SV Total RNA Isolation System (Promega, Madison, WI, USA). A thousand ng of total RNAs were used for cDNA synthesis using the SMART MMLV Reverse Transcriptase (Takara Bio, Mountain View, CA, USA). For RT-PCR, one microliter of cDNA was used as the template, and a 348-bp FoMV CP fragment was amplified using GoTaq DNA polymerase (Promega, Madison, WI, USA). The cDNA was diluted 40 times and then 2 ul were used for the RT-qPCR. The RT-qPCR was done using SsoAdvanced Universal SYBR Green Supermix (Bio-Rad, Hercules, CA, USA) as instructed. The *N. benthamiana* Actin2 gene and Z. mays Actin gene were used as the internal control. The relative gene expression was quantified using the delta-delta Ct method. A C1000 and a CFX384 thermocycler were used for RT-PCR and RT-qPCR (Bio-Rad, Hercules, CA, USA), respectively. RT-PCR and RT-qPCR primer sequences are listed in Supplemental Table 1.

## 3. Results

### 3.1. The DAG motif and its variants are commonly present in the N-terminus of potyvirus coat protein

To gain insights into setting up the conditional transmission of FoMV by introducing the potyvirus CP DAG motif, we first compared the CP amino acid sequence of different potexviruses and potyviruses. The alignment showed that both the N-terminus of potexvirus and potyvirus CP are highly variable (Fig. 1A & 1B). The C-terminus of the potexvirus CP is also not conservative, while the C-terminus of the potyvirus CP is comparably conserved. For the potexvirus, the DAG motif was only found in the N-terminus of PAMV CP at the position of 14-16 (Fig. 1A). While a single or double DAG motif, or its variants, was found in the N-terminus of all the aligned potyvirus CPs (Fig. 1B). The DAG motifs were usually located near the N-terminus of the CPs, within the range of their N-terminus 15 amino acid residues. This further confirms the DAG motif (or its variant) is indispensable for the aphid transmissibility of potyviruses, thus we sought to set up the conditional transmission of FoMV by introducing the DAG motif to the N-terminus of its CP.

**Figure 1:**
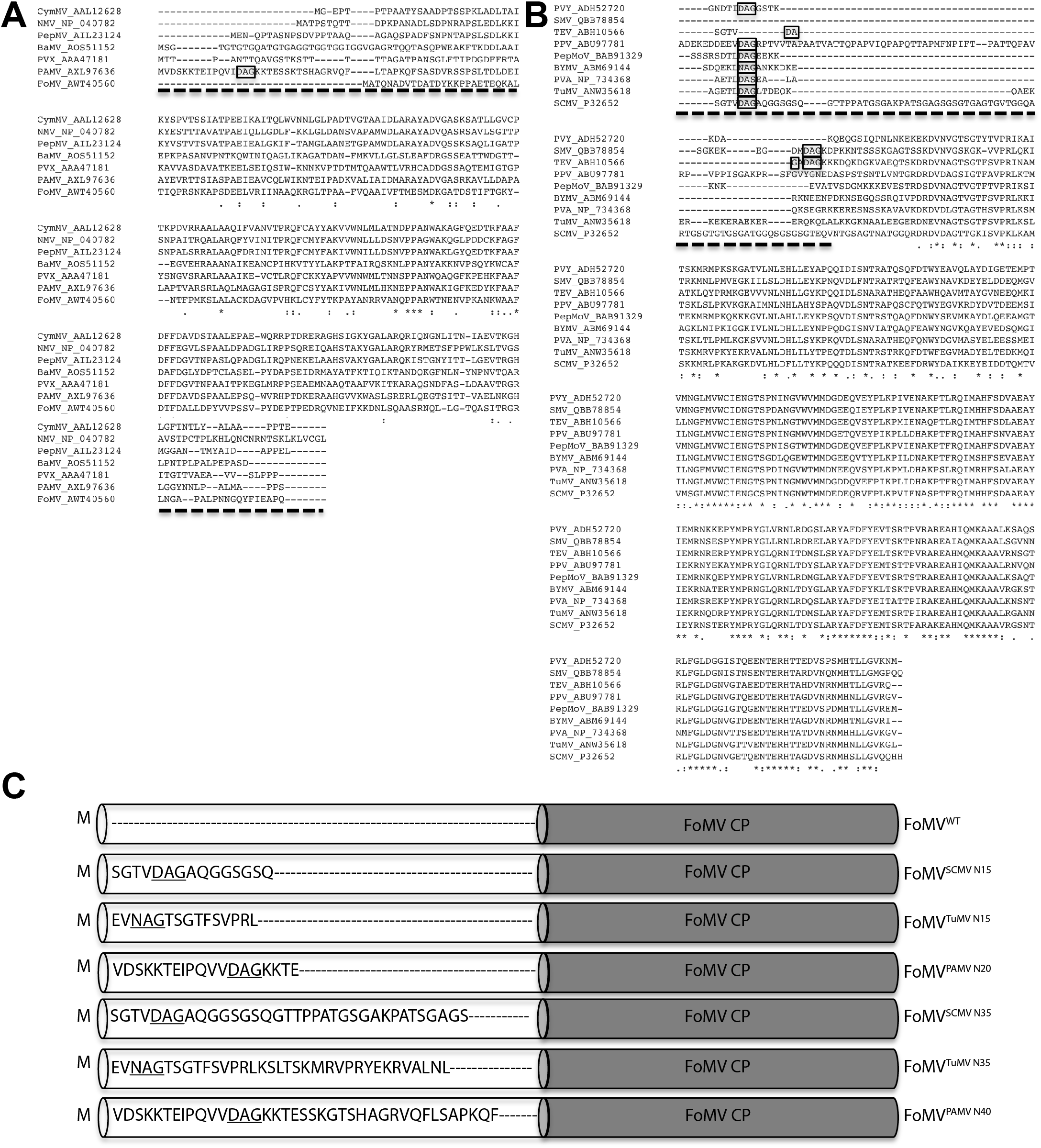
Potexvirus and potyvirus coat protein amino acid sequence alignment and the mutants constructed in this research. The coat protein amino acid sequence of several Potexviruses (A) and potyviruses (B) are aligned. CymMV, *Cymbidium mosaic potexvirus*; NMV, *Narcissus mosaic virus*; PepMV, *Pepino mosaic virus*; BaMV, *Bamboo mosaic virus*; PVX, *Potato virus x*; PAMV, *Potato aucuba mosaic potexvirus*; FoMV, *Foxtail mosaic virus*; PVY, *Potato virus y*; SMV, *Soybean mosaic virus*; TEV, *Tobacco etch virus*; PPV, *Plum pox virus*; PepMoV, *Pepper mottle virus*; BYMV, *Bean yellow mosaic virus*; PVA, *Potato virus A*; TuMV, *Turnip mosaic virus*; SCMV, *Sugarcane mosaic virus*. The accession number of each sequence is listed after the virus abbreviation. The dashed underlines highlight the unconserved N- and C-terminus. The boxes indicate the DAG motif or its variants. Identical amino acid residues that are highly conserved are highlighted by stars, and similar amino acid residues are indicated by dots. FoMV mutants constructed in this research are shown in C, with the amino acid residues of the introduced N-terminal tail are shown.

### 3.2. The recombinant FoMV viruses form infectious virus particles

Based on the sequence alignment information, we first introduced the potyvirus DAG motif into the FoMV CP by substituting amino acid residues within its N-terminus. In this case, we maintained the authentic full-length sequence of the FoMV CP, changing only a few amino acid residues, and expected these mutants would still be capable of forming infectious virus particles. Five mutants, FoMV^V8A-T9G^, FoMV^T12G^, FoMV^Y14A-K15G^, FoMV^V8A-T9G-T12G^, and FoMV^A6V-V8A-T9G^ were constructed (see supplemental Table 2). In the mutants FoMV^V8A-T9G^, FoMV^T12G^, and FoMV^Y14A-K15G^, a single DAG motif was introduced at positions 7-9, 10-12, and 13-15 within the N-terminus of FoMV CP, respectively. For mutant FoMV^V8A-T9G-T12G^, double DAG motifs were introduced at positions 7-9 and 10-12. For mutant FoMV^A6V-V8A-T9G^, a DAG motif was introduced at position 7-9, and the alanine residue at position six was substituted with valine. This mutant was made because we found an alanine residue that preceded the DAG motif quite often in the N-terminus of potyvirus CPs. The systemic infection of the WT and all five mutated FoMV was detected by RT-PCR at 5 days post agroinfiltration (data not shown). However, none of the mutated viruses that carry the DAG motif were aphid transmissible in the presence of TuMV HC-Pro protein (data not shown).

We then explored an alternative strategy where the N-terminal part of SCMV’s, TuMV’s, or PAMV’s CP, which contains the DAG motif, was added to the N-terminus of the full-length FoMV CP (Fig. 1C). The addition of N-terminal 15 or 35 amino acid residues of SCMV and TuMV CP, resulting in the recombinant virus termed FoMV^SCMV N15^, FoMV^TuMV N15^, FoMV^SCMV N35^, and FoMV^TuMV N35^. The addition of the N-terminal 20 and 40 amino acid residues to the N-terminus of FoMV CP are named FoMV^PAMV N20^ and FoMV^PAMV N40^ thereafter, respectively. First, we wanted to check if these recombinant viruses still can replicate. As shown in Fig. 2A, the WT FoMV viral RNA (vRNA) accumulation reached the highest level in inoculated *N. benthamiana* leaves at 5 days post agroinfiltration. The vRNA accumulation for the recombinant viruses was then compared in *N. benthamiana* leaves 5 days after agroinfiltration. We found all the recombinant viruses were still replicable, although the replication level was about 20-40% of the WT virus (Fig. 2B). The systemically infected leaves of the above agroinfiltrated *N. benthamiana* were then checked by RT-PCR at 5 days post agroinfiltration. The recombinant virus FoMV^SCMV N15^, FoMV^TuMV N15^, FoMV^PAMV N20^, FoMV^SCMV N35,^ and FoMV^PAMV N40^ were able to establish a systemic infection in *N. benthamiana* (Fig. 2C, upper panel). Systemically infected *N. benthamiana* leaf tissues were used as the source tissues for rub inoculation into *Zea mays* cv. Golden Bantam. All FoMV recombinants, except FoMV^TuMV N35^ could also infect *Z. mays* systemically at 2 weeks after rub inoculation (Fig. 2C, lower panel). Next, virus particles were purified from systemically infected *N. benthamiana* leaf tissues, and the purified virus particles were observed under TEM. As expected, the above mutated virus FoMV^SCMV N15^, FoMV^TuMV N15^, FoMV^PAMV N20^, FoMV^SCMV N35^, and FoMV^PAMV N40^ were able to form virus particle. This means the addition of exogenous N-terminal tails does not affect the assembly of the virus particles, which is a prerequisite for aphid-mediated virus transmission. For aphid transmission experiments , the recombinant virus FoMV^TuMV N35^ was not considered as it could not establish virus systemic infection, probably because this recombinant virus could not form infectious virus particles or lose the movement ability.

**Figure 2:**
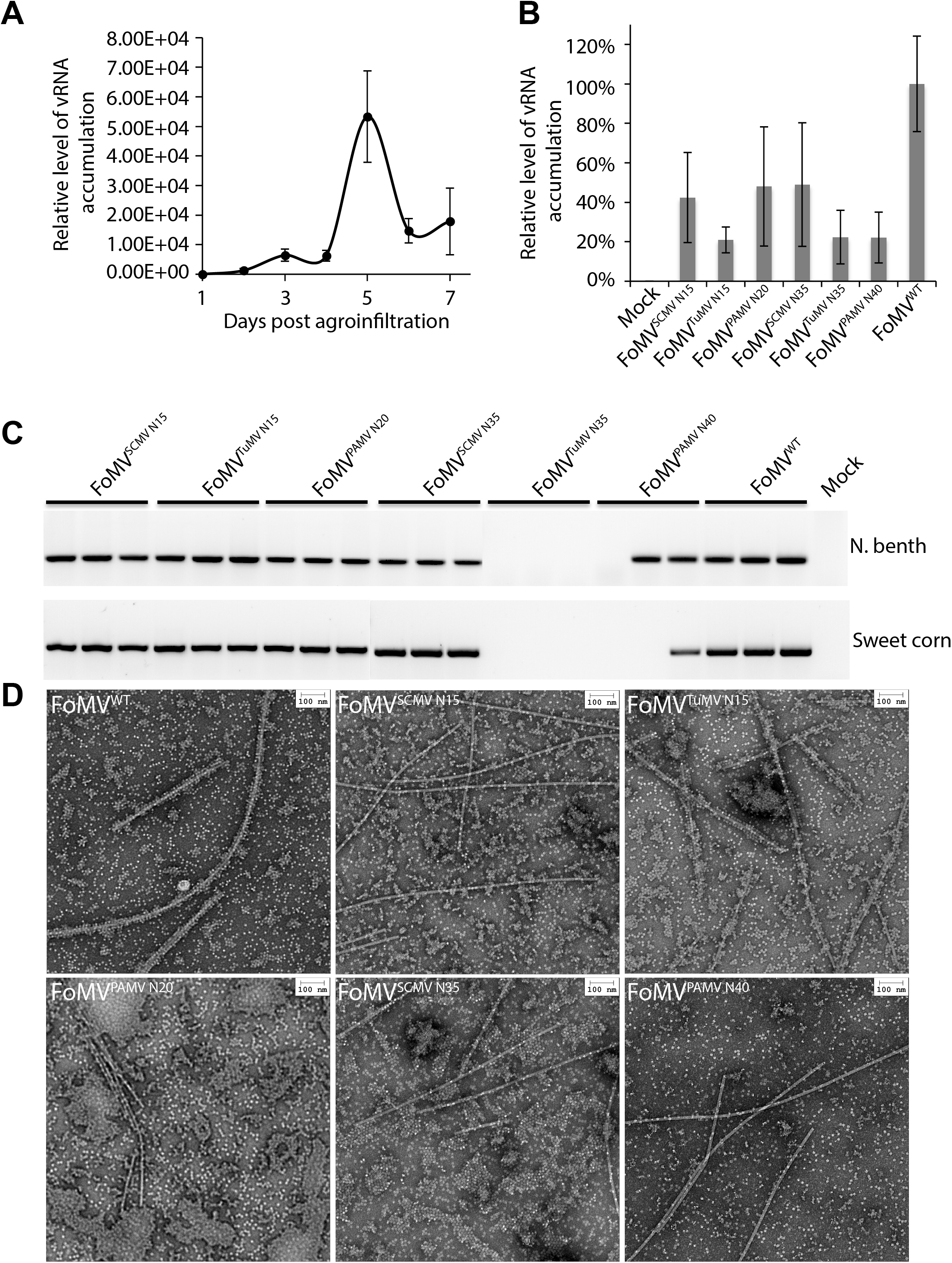
FoMV mutant characterization. *N. benthamiana* leaf tissues were agroinfiltrated with WT FoMV, and the accumulation of vRNA was quantified by RT-qPCR at different days up to 7 days (A). The FoMV mutant agroinfiltrated *N. benthamiana* tissues were collected at 5 days after agroinfiltration, and the tissues were processed for RT-qPCR (B). Mean values ± standard deviations (SD) from three independent experiments are shown (A & B). The infection of FoMV mutant systemic infected *N. benthamiana* (C, upper panel) or *Z. mays* (C, lower panel) leaf tissues were checked by RT-PCR. Virions were purified from the systemic infected *N. benthamiana* leaf tissues and observed with transmission electron microscopy (D).

### 3.3. The recombinant viruses, in particular the FoMV^SCMV N35^, are aphid transmissible

Next, we then explored if the recombinant FoMV viruses are aphid transmissible. *N. benthamiana* leaf tissues co-infected with the above FoMV recombinant viruses and TuMV/6K2:GFP. The co-infected plants were then used as the source tissue for virus transmission (Fig. 3A). *N. benthamiana* co-infected with TuMV/6K2:GFP and WT FoMV or with the recombinant FoMV’s, were used as the negative control. As shown in Fig. 3B, the recombinant virus FoMV^SCMV N15^, FoMV^PAMV N20^, FoMV^SCMV N35^, and FoMV^PAMV N40^, could be transmitted to *N. benthamiana* efficiently by aphids However, only the recombinant virus FoMV^SCMV N35^ could be transmited to *Z. mays* efficiently (Fig. 3C). On average, the transmission efficiency ranged from 30-80% for *N. benthamiana* and 10-75% for *Z. mays* (Fig. 3D). The negative controls did not show any virus transmission (data not shown). In conclusion, these data suggest the recombinant viruses, in particular the FoMV^SCMV N35^, are aphid transmissible in the presence of TuMV HC-Pro protein.

**Figure 3:**
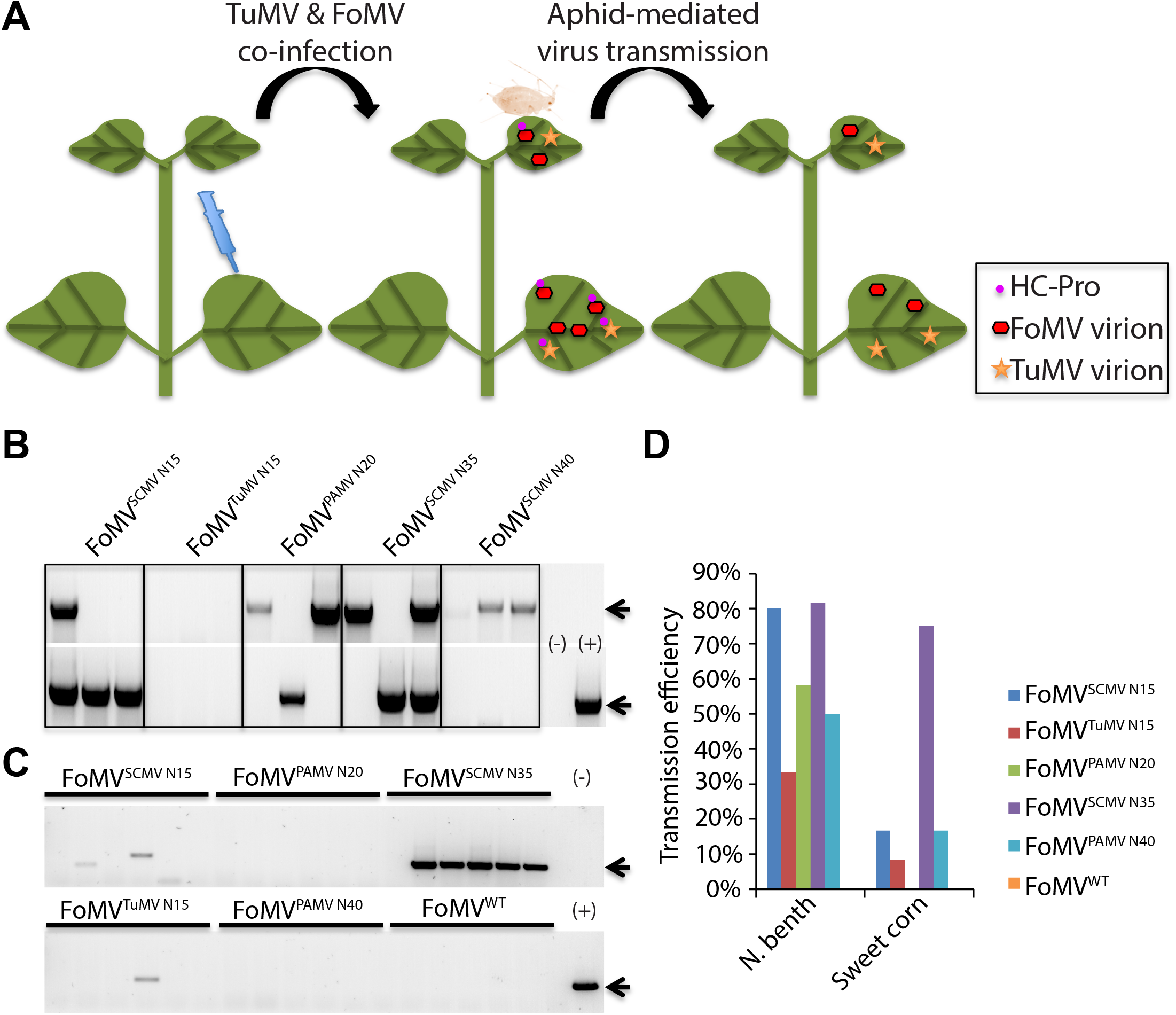
FoMV aphid transmission. Fig. 3A shows how the aphid transmission was performed in the presence of TuMV/6K2:GFP co-infection. *N. benthamiana* leaf tissues co-infected with TuMV/6K2:GFP and FoMV recombinant virus were used as the source tissues for aphid transmission. The aphids were starved for 4 h, and fed on the source tissue for 5 min before transferred to the healthy plants. One representative aphid transmission from infected *N. benthamiana* to healthy *N. benthamiana* (B) or healthy *Z. mays* (C) is shown. (B and C) The presence of virus infection was confirmed by RT-PCR. Upper panels: the amplification of a 348bp fragment of FoMV; Lower panels: the amplification of internal control; (+): the plasmid pCambia1380/FoMV used as the PCR template. The data from two times experiments are shown as (D). The value of how many plants were infected out of the tested plants are shown.

## 4. Discussion

The naturally occurring DAG motif of the potexvirus PAMV can make PAMV and other non-aphid-transmissible potexviruses, such as PVX, aphid-transmissible in mixed infections with a potyvirus (Baulcombe et al., 1993). We demonstrated that the N-terminal portion of different potyvirus CPs, which contains the DAG motif, can also be used to engineer aphid transmission for potexviruses. The N-terminal portion of the CP from the aphid-transmissible potyviruses, TuMV or SCMV, or from the previously studied DAG containing potexvirus (PAMV), made FoMV aphid transmissible when co-infected with a potyvirus (Fig. 3). The presence of the potyvirus protein HC-Pro is required as the transmission can only happen during co-infection with a potyvirus.

The importance of the context of the CP DAG motif in virus aphid transmission has been demonstrated (López-Moya et al., 1999). The presence of the DAG motif does not guarantee transmissibility, and the context in which the DAG or equivalent motif is found plays a role in the process. This is consistent with what we have found, simply introducing the DAG motif in the N-terminus of FoMV CP by substituting certain amino acids did not make FoMV aphid transmissible. The virus transmission could happen only when part of the N-terminal tail of potyvirus CPs were added to the N-terminus of FoMV CP (Fig. 3). We also noticed that the DAG motifs from different viruses showed varied transmission efficiency in the presence of the same HC-Pro protein (Fig. 3B & 3C). Our sequence alignment showed that the N-terminus of potyvirus CP is highly variable, although the DAG motif is commonly present near the N-terminus of the CP (Fig. 1B). These indicate the amino acid residues surrounding the DAG motif are important for the virus aphid transmission, probably by affecting the HC-Pro accessibility to the DAG motif.

Overall, our data shows the aphid transmission was more successful to the model plant *N. benthamiana* compared to *Z. mays*, where transmission efficiency varied from 30% to over 80%, depending on the N-terminus addition. We also noticed that one recombinant virus, FoMV^PAMV N20^, could only be transmitted by aphids to *N. benthamiana* plant but not to *Z. mays* (Fig. 3D). Similarly, the recombinant virus, FoMV^SCMV N15^, could be transmitted to *N. benthamiana* plant efficiently, but this was not the case when it was transmitted to *Z. mays* (Fig. 3D). The highest aphid transmission efficiency was observed for FoMV^SCMV N35^ for both host plants (Fig. 3). The DAG motif may play dual functions by mediating aphid transmission and virus movement for some viruses. In the case of *Zucchini yellow mosaic virus* (ZYMV), the N-terminal tail of its CP doesn't seem important for the virus infection as the virus is still infectious when the N-terminal tail 43 amino acid residues are deleted (Arazi et al., 2001). However, for the *Tobacco vein mottling virus* (TVMV), the DAG motif is critical for the virus systemic infection (López-Moya and Pirone, 1998). In this previous study mutation of the DAG motif abolished the virus cell-to-cell movement, but not the vRNA replication in the cells. Thus, it will be interesting to investigate whether the higher transmission efficiency of the recombinant virus, FoMV^SCMV N35^, to *Z. mays*, is due to a better fitness of this virus in sweet corn, which is a susceptible host of SCMV infection.

Although we have established engineering aphid transmission of FoMV can be successful, our results demonstrate not all DAG motifs are equal and additional work is needed. In our study, only one aphid species (*M. persicae*) and one HC-Pro (TuMV) were used. It is known that the HC-Pro protein of Watermelon mosaic virus (WMV) can assist the aphid transmission of TuMV, while the HC-Pro protein of TuMV is ineffective for the transmission of WMV (Sako and Ogata, 1981). Also, different aphid species can transmit the same virus at different efficiency, in the case of ZYMV and TuMV (Dombrovsky et al., 2005). Efforts are still needed to figure out the best combination of HC-Pro source and aphid species for a particular aphid-mediated plant virus transmission. It is also not known whether this strategy can be applied to other non-aphid transmissible plant viruses, such as viruses with icosahedral virions, of which the nucleic acid encapsidation capacity is limited, so the additional amino acids may influence virus particle assembly. The strategy that we have established here only applies to aphid-mediated virus transmission. Thus, it is necessary to explore the other applicable strategies for the conditional transmission of plant viruses.

## Author Contributions

CLC conceived the original research plans; JJ and EY performed 565 the experiments; JJ analyzed the data; CLC and JJ supervised the experiments; JJ and CLC wrote the article with contributions of all the authors; CLC agrees to serve as the contact author responsible for communication and distribution of samples

## Acknowledgments

We thank Professor Steven A. Whitham for kindly providing the pCambia1380/FoMV infectious clone. We thank Professor Jean-François Laliberté for the pCambia0380/TuMV:6K2-GFP infectious clone. This research was supported by Defense Advanced Research Projects Agency (DARPA) agreement HR0011-17-2-0053 t and by a US National Science Foundation award award 1723926 to CLC. The views and conclusions contained in this document are those of the authors and should not be interpreted as representing the official policies, either expressed or implied, of DARPA or the U.S. Government.

---

**Supplemental Table 1: Primers used in this study.**

**Supplemental Table 2: Additional mutants constructed in this research.**

