## Supplemental Table 1 for "Aphid transmission of a Potexvirus, *Foxtail mosaic virus*, in the presence of the Potyvirus helper component proteinase"

Supplemental Table 1: Primers used in this study

| **Purpose** | **Primer** | **Sequence (5’→3’)** |
| --- | --- | --- |
| pCambia1380/FoMV^SCMV N15^ | F1 | AACCAACATCAGTGAAGAGAAACCCTTAGGAGAGTTAACAACGGGCCCC |
|  | R1 | CATTTTGTGTTGCTTGGCTTCCACTGCCGCCTTGTGCACCCGCATCAACAGTTCCCGACATTGTGTCCTGAAATGATGA |
|  | F2 | CAGTGGAAGCCAAGCAACACAAAATGCCGAC |
|  | R2 | ATACTACTGATCATTAATTAAGGGTCTAGATTACTGAGGTGCCTCGATG |
| pCambia1380/FoMV^TuMV N15^ | R1 | CATTTTGTGTTGCGAGTCTGGGTACACTGAAAGTTCCAGAGGTTCCAGCGTTTACTTCCATTGTGTCCTGAAATGATGAGGT |
|  | F2 | TGTACCCAGACTCGCAACACAAAATGCCGAC |
| pCambia1380/FoMV^PAMV N20^ | R1 | CATTTTGTGTTGCTTCAGTCTTCTTGCCGGCGTCAACCACTTGAGGAATTTCCGTTTTCTTAGAATCAACCATTGTGTCCTGAAATGATGAGGTCACA |
|  | F2 | CAAGAAGACTGAAGCAACACAAAATGCCGAC |
| pCambia1380/FoMV^SCMV N35^ | R1 | CATTTTGTGTTGCAGATCCTGCCCCTGAGGTGGCTGGTTTTGCTCCACTACCTGTTGCTGGTGGTGTTGTCCCTTGGCTTCCACTGCCGCCTTGTGCA |
|  | F2 | AGGGGCAGGATCTGCAACACAAAATGCCGAC |
| pCambia1380/FoMV^TuMV N35^ | R1 | CATTTTGTGTTGCGAGGTTTAGAGCCACTCTTTTCTCGTATCTTGGCACGCGCATCTTGCTTGTCAGACTCTTGAGTCTGGGTACACTGAAAGTTCCA |
|  | F2 | GGCTCTAAACCTCGCAACACAAAATGCCGAC |
| pCambia1380/FoMV^PAMV N40^ | R1 | CATTTTGTGTTGCGAACTGCTTAGGGGCAGAGAGGAATTGAACGCGTCCAGCATGTGAGGTTCCTTTTGAACTTTCAGTCTTCTTGCCGGCGTCAACCA |
|  | F2 | CCCTAAGCAGTTCGCAACACAAAATGCCGAC |
| RT-qPCR | Actin2F | GGTAACATTGTGCTCAGTGGTGG |
|  | Actin2R | AACGACCTTAATCTTCATGCTGC |
|  | FoMV-RT-qPCR-F | TCTGTACCGTACGATGAGCCC |
|  | FoMV-RT-qPCR-R | GTTGAGTCTCCCGCGCGTGAT |
| RT-PCR | FoMV-RT-PCR-F | CAGAAGGCACTCACCATTCA |
|  | FoMV-RT-PCR-R | CTTGTTAGCTTTGGGCACATTC |

Note: The primer F1 and R2 to make construct pCambia1380/FoMV^TuMV N15^, pCambia1380/FoMV^PAMV N20^, pCambia1380/FoMV^SCMV N35^, pCambia1380/FoMV^TuMV N35^ and pCambia1380/FoMV^PAMV N40^ is the same as the F1 and R2 primer for pCambia1380/FoMV^SCMV N15^.
