## Supplementary figures and images for "Aphid transmission of a Potexvirus, *Foxtail mosaic virus*, in the presence of the Potyvirus helper component proteinase"

### Supplemental Table 2

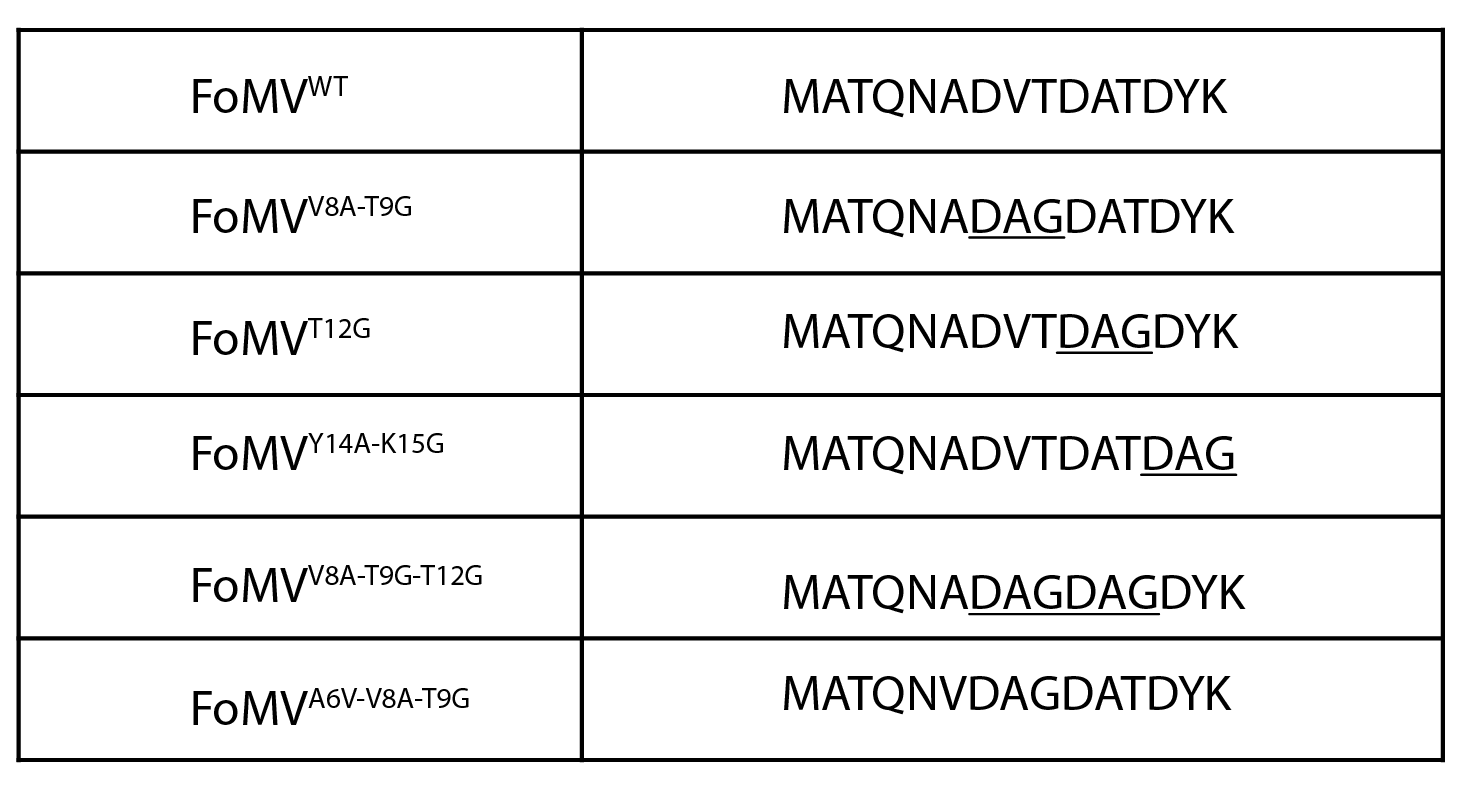
